# Patterns of siderophore production and utilization at Station ALOHA from the surface to mesopelagic waters

**DOI:** 10.1101/2022.10.04.510528

**Authors:** Randelle M. Bundy, Lauren E. Manck, Daniel J. Repeta, Matthew J. Church, Nicholas J. Hawco, Rene M. Boiteau, Jiwoon Park, Edward F. DeLong, Mak A. Saito

## Abstract

The North Pacific subtropical gyre is a globally important contributor to carbon uptake and an oligotrophic ecosystem primarily limited by nitrogen. The microbial community is also seasonally exposed to low iron due to biological consumption and seasonally variable iron delivery. In this study, we examined changes in iron uptake rates, dissolved siderophore concentrations, and siderophore biosynthesis at Station ALOHA across time (2013-2016) and depth (surface to 500 m) to observe changes in iron acquisition and internal cycling by the microbial community. The genetic potential for siderophore biosynthesis was widespread throughout the upper water column, and biosynthetic gene clusters peaked in spring and summer along with siderophore concentrations, suggesting changes in nutrient delivery, primary production, and carbon export impact iron acquisition over the seasonal cycle. Dissolved iron turnover times, calculated from iron-amended experiments conducted using surface (15 m) and mesopelagic (300 m) waters, ranged from 9-252 days. The shortest average turnover times at both depths were associated with inorganic iron additions (14±9 days) and the longest with iron bound to strong siderophores (148±225 days). Uptake rates of siderophore-bound iron were faster in the mesopelagic waters than in the surface, leading to high Fe:C uptake ratios of heterotrophic bacteria in the upper mesopelagic. The rapid cycling and high demand for Fe at 300 m suggests differences in microbial metabolism and iron acquisition in the mesopelagic compared to surface waters. Together, changes in siderophore production and consumption over the seasonal cycle suggest organic carbon availability impacts iron cycling at Station ALOHA.

**Scientific Significance Statement:** Microbial community production in the subtropical oligotrophic North Pacific is limited by macronutrients such as nitrogen. However, dissolved iron is another important micronutrient that has seasonal inputs from dust and passing eddies, keeping the availability of iron low and episodic. Little attention has been paid to the microbial strategies for dealing with low iron to support primary production in the oligotrophic ocean, or how limited iron availability impacts the processing of sinking particulate organic carbon in this region. In this study, we explore iron cycling including siderophore production and uptake by the microbial community throughout the water column at Station ALOHA to examine how the microbial community adapts and responds to changing iron and carbon availability on seasonal timescales.

## 1. Introduction

Oligotrophic subtropical gyres comprise the world’s largest ocean biome (Longhurst 2010). Microbial communities inhabiting the oligotrophic ocean are exposed to low concentrations of nitrogen, phosphorous (Karl et al. 1997; Fitzsimmons and Boyle 2014; Letelier et al. 2019), and dissolved iron (Fitzsimmons et al. 2015). Long-term studies by the Hawaii Ocean Time-series (HOT) at Station ALOHA in the North Pacific subtropical gyre demonstrate how seasonal to interannual-scale changes in nutrient supply can impact microbial communities and carbon export (Karl and Lukas 1996; Letelier et al. 2019). The resulting time series captures both the relatively stable conditions of the oligotrophic ocean and the importance of episodic and seasonal-scale delivery of nutrients to the upper ocean from storms, eddies, dust inputs, and the shoaling and deepening of the pycnocline (DiTullio and Laws 1991; Fitzsimmons et al. 2015; Hayes et al. 2015; Pinedo-González et al. 2020; Hawco et al. 2022). Although this ecosystem is characterized by perennial nitrogen limitation, the microbial community of the euphotic zone is also consistently exposed to low dissolved iron (dFe) concentrations, particularly in the deep chlorophyll maximum (DCM; (Hogle et al. 2022, Hawco et al. 2022)), and on sub-decadal timescales, control of primary production as supported by nitrogen (N_2_) fixation is thought to oscillate between phosphate and iron availability (Letelier et al. 2019).

In seawater, a majority of dFe is complexed by a pool of organic ligands (FeL) that have a high diversity of chemical structures. Complexation by organic ligands allows dFe to accumulate to concentrations above that set by its inorganic solubility (Gledhill and Buck 2012) and may help to retain Fe within the euphotic zone where it is needed to support primary production (Tortell et al. 1999). However, ligands significantly alter the bioavailability of the dFe pool by sequestering Fe in organic complexes that require specific cellular transport systems in order to be accessed (Sutak et al. 2020). Changes in the external supply of Fe to the upper ocean at Station ALOHA have been shown to trigger organic ligand production on the timescale of days (Fitzsimmons et al. 2015).

Marine microorganisms, including both photoautotrophic and heterotrophic bacteria, have significant cellular Fe requirements and thus are equipped with molecular strategies for acquiring Fe from the Fe-scarce marine environment (Hopkinson et al. 2005; Sutak et al. 2020). A class of the strongest organic Fe-binding ligands, called siderophores, are secreted by marine bacteria and fungi in response to Fe deficiency as a high-affinity uptake strategy to solubilize and stabilize Fe in a molecular form that can be taken up by dedicated membrane transporters (Sandy and Butler 2009; Vraspir and Butler 2009). Many structurally diverse siderophores have been identified, and they are biosynthesized by two classes of enzymatic pathways; non-ribosomal peptide synthetase (NRPS) pathways and/or NRPS-independent siderophore (NIS) synthetase pathways (Sandy and Butler 2009; Hider and Kong 2010). Siderophore biosynthesis is regulated by Fe concentrations, where elevated Fe suppresses synthesis (Sandy and Butler 2009). Some microbes can both synthesize and internalize siderophores, while others only possess outer membrane receptors for uptake of exogenous siderophores that they cannot produce (Cordero et al. 2012; Kramer et al. 2020). Internalizing siderophore-bound Fe requires either specific transport pathways or the ability to extracellularly reduce Fe(III) to Fe(II), releasing it from the siderophore complex; thus Fe complexed to siderophores has low bioavailability to the microbial community as whole (Shaked and Lis 2012), but is readily bioavailable to microbes that possess the relevant transport systems.

Dissolved siderophores represent 2-10% of the total dissolved Fe-binding ligand pool present at Station ALOHA, and they likely play an important role in the competition for Fe and in solubilization of Fe from particles (Bundy et al. 2018). North of Station ALOHA in the North Pacific transition zone, where Fe inputs from dust in the spring are elevated (Pinedo-González et al. 2020), siderophore concentrations and the transcription of siderophore biosynthesis and uptake genes are also elevated, further connecting the production of siderophores as a mechanism for solubilizing Fe from particles (Park et al. 2023). The metabolic potential for siderophore production and uptake is prevalent in the North Pacific subtropical gyre, transition zone, and the subpolar high nutrient low chlorophyll region (Hogle et al. 2022; Park et al. 2023).

Although the importance of siderophore production as a microbial Fe acquisition strategy in the marine environment has been recognized for some time (Mawji et al. 2008; Boiteau et al. 2016, 2019; Bundy et al. 2018; Moore et al. 2021; Manck et al. 2022; Park et al. 2023), little is known about their seasonal variability or turnover times. Understanding the biosynthesis and turnover times of siderophore-bound Fe will be important for interpreting their bioavailability and influence on Fe recycling and retention in the upper ocean. To address these gaps in our understanding of siderophore biogeochemistry in the marine environment, we explored changes in siderophore distributions, biosynthesis potential and turnover times at Station ALOHA across time and depth. By placing this study in the rich context of biogeochemical measurements made by the HOT program, we aim to better connect siderophore production and uptake to environmental conditions, in an effort to understand the impact of siderophores on microbial Fe acquisition and biogeochemical cycling in this important and well-studied oligotrophic ecosystem.

## 2. Methods

### 2.1 Radioactive iron uptake and siderophore bioavailability experiments

Fe uptake experiments were performed at Station ALOHA (Fig. 1A) on two expeditions (HOT278 in November 2015 and KM1605 in March 2016) at 15 m and 300 m. Four treatments were used in all experiments: FeCl_3_ (denoted as “FeL”), ferrioxamine B, ferrioxamine E, and Fe-amphibactin. Experiments at 15 m were performed both in the light and dark, and experiments at 300 m were performed only in the dark. Fe uptake was measured in the >2.0 µm and 0.2-2.0 µm size fraction after 12 hours. 500 nmol L^-1^ of nitrate was added to each of the 15 m experiments to alleviate possible nitrogen limitation, which might have restricted Fe uptake (Table S1).

**Figure 1.**
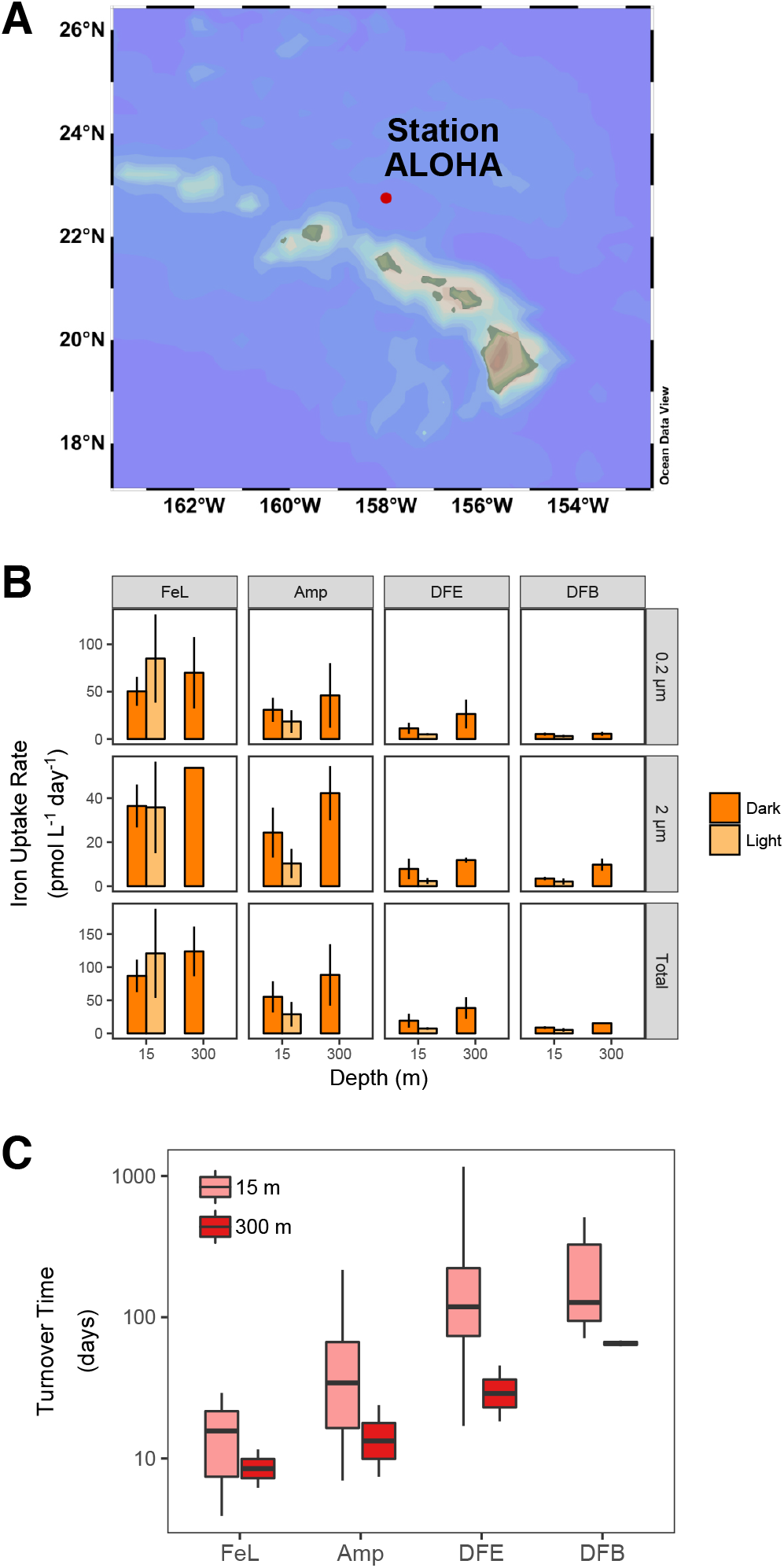
**(A)** Map of the study region including the location of Station ALOHA (red dot). **(B)** Dissolved iron uptake rates (pmol L^-1^ day^-1^) in the surface (15 m) and mesopelagic (300 m) at Station ALOHA for naturally complexed iron (FeL) and iron bound to the siderophores amphibactin (Amp), ferrioxamine E (DFE), and ferrioxamine B (DFB). Dark and light measurements are displayed for both the 0.2-2.0 µm and >2.0 µm size fraction. Total rates in the bottom panels represent the sum of both size fractions for each treatment. Mean values from all measurements for a given treatment are displayed and error bars represent the standard error of the mean. (B) Turnover time (days) of iron in each treatment at 15 m and 300 m based on rates of total iron uptake in the dark. Box plots display the interquartile range with median values plotted as a solid horizontal line. The y-axis is on a log_10_ scale. Turnover times at 300 m were statistically different from 15 m only in the DFB treatment (pairwise t-test, *p* < 0.05).

The isolation of amphibactin ligands used in this experiment has been described in detail elsewhere (Bundy et al. 2018), and the other ligand standards were commercially available. Radioactive Fe ligand stocks were prepared using a 1 mCi ^55^Fe stock (Perkin Elmer; Lot #091514) with a specific activity of 44.66 mCi mg^-1^ and an activity of 35.55 mCi mL^-1^. Before making the ligand stocks, the concentration of the ^55^Fe stock solution was calculated by converting the initial activity on the stock date to the current activity using the half-life of ^55^Fe (λ = 2.73 years) and the number of days since the stock date. A primary ^55^Fe stock of 55 µmol L^-1^ was used to prepare the ^55^Fe-ligand and ^55^FeCl_3_ stocks. The FeCl_3_ stock (FeL treatment) was prepared in pH 2 (Optima HCl, Fisher Scientific) type I deionized water (MilliQ) while the siderophore stocks were prepared in type I deionized water. Ligand stocks were equilibrated for at least 24 hours, but up to 5 days, before use, and ligands were present in three times excess of ^55^Fe. Duplicate ligand stocks were made with the stable isotope ^57^Fe (Cambridge Isotopes) to set up replicate treatments for cell counts that did not contain radioactivity.

Prior to beginning experiments at sea, quench curves were made to account for physical and chemical quench. Physical quench was accounted for by placing a blank filter in 10 mL of scintillation cocktail in triplicate and counting for 10 minutes on the liquid scintillation counter. Physical quench was also assessed by filtering a range of seawater volumes (10-500 mL) onto separate 0.2 µm filters containing no ^55^Fe, and then placing each filter into 10 mL of scintillation cocktail and counting for 10 minutes. These physical quench controls account for whether the presence of cells or the filter itself impacts the counting efficiency of the liquid scintillation counter. The physical quench was found to be extremely small (< 1 % of blanks), and for all further analyses, blank filters were used for quench correction. Chemical quench was assessed by setting up 10 separate vials of scintillation cocktail, each with the same activity of ^55^Fe and increasing acetonitrile additions (0.1-10 nmol L^-1^). The chemical quench was also found to be very small (< 5% of the blank) and was incorporated into the counting efficiency of the method for all samples.

To set up the incubation experiments, trace metal clean seawater collected from X-Niskin bottles (Ocean Test Equipment) from 15 m and 300 m was dispensed into polycarbonate bottles. Four of the bottles were spiked with 1 nmol L^-1^ of the ^57^Fe ligand stocks and the rest were spiked in duplicate with 1 nmol L^-1^ of the ^55^Fe ligand stocks. A final concentration of 0.01% glutaraldehyde was added to four of the bottles to serve as “dead controls” and were incubated for 30 minutes before the experiment began to account for non-specific ^55^Fe adsorption. All bottles were incubated in surface seawater flow-through incubators and dark treatments were double-bagged in black polyethylene bags. After 12 hours, a flow cytometry sample was taken from each of the ^57^Fe treatments and was preserved with 0.01% glutaraldehyde for 30 minutes before being flash-frozen in liquid nitrogen and stored at −80°C. At the end of the 12 hour experiment, each bottle was sequentially filtered onto acid washed 2.0 µm and 0.2 µm filters using low pressure vacuum filtration. Both filters were washed three times with an oxalate wash to remove extracellular Fe (Tovar-Sanchez 2003).

A key assumption of these experiments is that all Fe remains complexed to the respective added siderophore throughout the duration of the experiment (12 hours). To confirm this was the case, the remainder of the treatments spiked with the ^57^Fe-siderophore stocks were filtered with a 0.2 µm polycarbonate filter, and the filtrate was pre-concentrated onto a solid phase extraction column (Bond Elut ENV, Agilent Technologies) and were treated and analyzed in the same manner as the dissolved siderophore samples from the water column as described below. ^57^Fe peaks from LC-ICPMS analyses were examined to determine whether any free ^57^Fe not associated with each of the added siderophores was present. No significant free ^57^Fe was observed in these treatments, confirming the full complexation of each ligand in the siderophore treatments (data not shown).

Radioactive decay on each filter and in each treatment was determined using a liquid scintillation counter (Beckman-Coulter LS6500). ^55^Fe disintegrations per minute (dpm) were background and dead control corrected and converted to a concentration of Fe uptake using a standard curve. Fe uptake rates were then calculated based on the total amount of Fe that was incorporated into biomass onto each filter size fraction and in each treatment. Total Fe uptake over the 12-hour incubation was converted to total Fe uptake per day (over 24 hours). We then estimated Fe uptake rates for heterotrophic bacteria specifically, which we assumed to be the dominant community responsible for any observed Fe uptake in the dark (Table S1). The Fe uptake per heterotrophic bacteria cell per day (amol Fe cell^-1^ day^-1^) was estimated using flow cytometry cell counts of heterotrophic bacteria (in the ^57^Fe treatments) and the total Fe uptake in the dark (Table S1). Fe:C ratios (μmol:mol) for heterotrophic bacteria were calculated using a carbon quota of 12.4 fg C cell^-1^ (Strzepek et al. 2005; Boyd et al. 2015) and assuming a growth rate of 1 day^-1^ (Jones et al. 1996, (Table S1)). As an additional constraint on Fe:C per cell, Fe uptake rates were compared to average rates of bacterial carbon production measured at or near Station ALOHA – 23±8 nmol C L^-1^ day^-1^ and 0.9±1.3 nmol C L^-1^ day^-1^ at 5 m and 300 m, respectively. The rate of bacterial production at 5 m is an average value of those measured at Station ALOHA (Viviani and Church 2017). The mean rate at 300 m was calculated from measurements conducted on three cruises near Station ALOHA in the North Pacific subtropical gyre – KOK1507, KM1605, and KM1709. All data are publicly available on the Simons Collaborative Marine Atlas Project. In each case, leucine incorporation was measured using the methods described by Smith and Azam (1992) and a conversion factor of 1.5 kg C mol^-1^ leucine was used to convert leucine incorporation to bacterial carbon production.

In addition to Fe uptake rates, the turnover rate of Fe in each treatment was calculated based on,

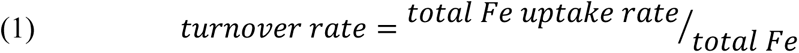

where the total Fe uptake rate (pmol L^-1^ day^-1^) is the sum of the radioactive Fe uptake in both size fractions and the total Fe is the radioactive Fe added (1 nmol L^-1^) plus the *in situ* dFe concentration (Bundy et al. 2018). The turnover time of Fe (in days) for each treatment was calculated as 1/turnover rate.

### 2.2 Environmental parameters

Data for concentrations of nitrate+nitrite (NO_3_^-^+NO_2_^-^), chlorophyll *a* (chl *a*), rates of primary production, and carbon flux at Station ALOHA were accessed for the study period, January 1, 2013-December 31, 2016. All data is publicly available via the HOT-Data Organization and Graphical System. Primary production is routinely estimated based on assimilation of ^14^C by the HOT program at depths of 5, 25, 45, 75, 100, and 125 m (Karl et al. 2021). Particulate carbon flux is measured from sediment traps at 150 m. Additional details on the sampling strategies used by the HOT program can be found elsewhere (Karl and Lukas 1996) and the analytical methods used for HOT measurements can be found online (https://hahana.soest.hawaii.edu/hot/methods/results.html). NO_3_^-^+NO_2_^-^concentrations were determined using the high-sensitivity chemiluminescent method (Dore and Karl 1996; Foreman et al. 2016). Depth integrated (0-150 m) fluxes and stocks were calculated using trapezoidal approximations, assuming homogenous mixing between 0-5 m. To integrate rates of primary production to 150 m, monthly mean rates measured at 150 m during the first 12 years of the HOT program (Karl et al. 2021) were added to the 0-125 m integrated value calculated during this study period. The *e*-ratio for each HOT cruise during the study period was calculated as the rate of particulate carbon flux measured at 150 m divided by the depth integrated (0-150 m) rate of primary production.

Because samples for dFe (< 0.2 µm) concentrations are not routinely measured by the HOT program, measurements made from previous studies at Station ALOHA (Fitzsimmons et al. 2015; Bundy et al. 2018) have been reproduced here. These measurements largely fall within the spring and summer months. Therefore, to visualize potential seasonal changes in dFe concentrations, 3-day average dFe concentrations at Station ALOHA during the study period were obtained from the MIT Darwin model (Dutkiewicz et al. 2015) via the Simons Collaborative Marine Atlas Project (CMAP): https://simonscmap.com/catalog/datasets/Darwin_Nutrient. Seasons were defined in this study as – Winter: December, January, February; Spring: March, April, May; Summer: June, July August; Fall: September, October, November. Additionally, vertical regions were defined here as: upper euphotic zone: ≤ 75 m, lower euphotic zone: > 75 m to ≤ 150 m, upper mesopelagic zone: > 150 m to ≤ 300 m, mid-mesopelagic zone: > 300 m to ≤ 500 m.

### 2.3 Siderophore collection and analyses

Dissolved siderophore samples were collected on six cruises from 2013-2016, each within 100 km of Station ALOHA (Table S2). Samples were collected via Teflon diaphragm pump or 8 L X-Niskin bottles on a trace metal rosette using a non-metallic line. Samples were filtered (0.2 µm Acropak 200 or Pall capsule) into acid-cleaned fluorinated high density polyethene 200 L barrels (expeditions from 2013-2014; Boiteau et al. 2013) or 20 L carboys (all other expeditions; Bundy et al. 2018). For each sample collected in 2013 and 2014, 20-800 L of filtered seawater was preconcentrated, and for all other samples 15-20 L of filtered seawater was collected. Filtered samples were preconcentrated at a flow rate of 15-18 mL min^-1^ onto 1000 mg solid phase extraction columns containing (Bond Elut ENV, Agilent Technologies). After preconcentration, each column was flushed with three column volumes of trace metal clean type I deionized water until dry and then was stored at −20°C until analysis.

Solid phase extraction columns were slowly thawed in the dark and then eluted with three column volumes of distilled or Optima grade methanol (Fisher Scientific). Methanol extracts were dried to approximately 0.5-1 mL using a SpeedVac (Thermo Fisher). Either 25 or 50 µL of the extract was first injected onto a PEEK-lined C8 column and compounds were separated using liquid chromatography (LC) on a Dionex 3000 high pressure liquid chromatography (HPLC) system. Details on the chromatography are presented in Boiteau et al. (2016) and Bundy et al. (2018). Briefly, compounds were separated using a 20 minute gradient of 95% solvent A/5% solvent B to 10% solvent A/90% solvent B (solvent A = 5 mM ammonium formate in type I deionized water, solvent B = 5 mM ammonium formate in distilled methanol), followed by an isocratic step from 90% to 95% solvent B (Boiteau et al. 2016; Bundy et al. 2018). Fe-bound compounds eluted from the column were monitored by inductively-coupled plasma mass spectrometry (ICP-MS) on an iCap-Q. The sample was introduced at a flow rate of approximately 50 µl min^-1^ and oxygen add gas was introduced at a flow rate of approximately 25 mL min^-1^ to minimize the deposition of reduced carbon on the cones. The ICP-MS was equipped with a perfluoroalkoxy micronebulizer (PFA-ST; Elemental Scientific), platinum sampler and skimmer cones, and a cyclonic spray chamber cooled to 0^◦^C. Counts per second of ^56^Fe, ^57^Fe, ^54^Fe and ^59^Co were monitored in kinetic energy discrimination mode (KED) and distinct peaks in ^56^Fe and ^57^Fe were integrated using in-house R scripts. Peak areas were converted to concentration using a four-point standard curve with ferrioxamine E, and were corrected for sensitivity changes throughout the run using the peak area of ^59^Co from a cyanocobalamin internal standard. All distinct quantifiable Fe peaks were categorized as putative siderophores and the concentrations of these peaks were summed for each sample to give a total siderophore concentration (Table S2).

To identify the putative siderophores, samples were then analyzed using LC coupled to electrospray ionization mass spectrometry (ESI-MS) on an Orbitrap Fusion (Thermo Scientific). The same chromatography was used, and the ICP-MS and ESI-MS data were aligned based on the retention time of the ^59^Co peak from cyanocobalamin in the ICP-MS trace and the presence of the cyanocobalamin *m*/*z* in the ESI-MS data (*m*/*z* = 678). Putative siderophores were identified using a targeted search of the MS^1^ *m*/*z* based on a database of known siderophores (Baars et al. 2014). Fragmentation data (MS^2^) was then examined and compared to literature values or in-silico fragmentation prediction (CFM-ID 3.0), and siderophores that had a matching MS^2^ fragment to literature values or in-silico prediction were assigned a structural identification (Table S2).

### 2.4 Siderophore biosynthetic gene cluster analysis

Details on the sampling, processing and data analyses of metagenomes collected at Station ALOHA are presented elsewhere (Mende et al. 2017; Luo et al. 2020). The ALOHA gene catalog consists of ∼8.9 million non-redundant genes detected from concentrated plankton biomass (0.22-1.6 μm) collected between the surface and 1,000 m over ∼1.5 years at near monthly frequency (Mende et al. 2017). To identify putative siderophore biosynthetic gene clusters (BGCs) within the ALOHA gene catalog, biomarker genes for both NRPS pathways and NIS pathways were first identified. To identify NIS pathways in the ALOHA gene catalog, hidden Markov models (HMMs) for the conserved domains pfam04183 (IucA/IucC siderophore synthases) and pfam06276 (FhuF-like reductases) were searched for in the ALOHA gene catalog using HMMER (Eddy 2011) which resulted in the detection of 20 NIS homologs (Table S3). NIS pathways are specific to siderophore biosynthesis, hence the 20 identified NIS homologs could be confidently attributed to siderophore biosynthesis. For NRPS biosynthetic pathways, we searched for the conserved domain pfam00501, an AMP-binding domain inclusive of the adenylation domain of NRPS modules. The resulting 28,000+ genes were then screened with antiSMASH (Medema et al. 2011) to determine the presence of a complete or near-complete NRPS module with adenylation, peptidyl carrier protein (pfam00550), and condensation (pfam00668) domains. This resulted in the detection of 132 NRPS homologs. Because NRPS biosynthetic pathways are not unique to siderophore biosynthesis, further examination of the identified NRPS homologs was required. Each NRPS homolog, along with neighboring genes from the ALOHA gene catalog contig assembly, were analyzed using the predictive tools within the antiSMASH pipeline with default parameters for the relaxed detection strictness setting. In addition to the default features, MIBiG cluster comparison and Cluster pfam analysis were also performed. This enabled predictions of the amino acid specificity for each adenylation domain within the NRPS modules, detection of additional conserved domains, and BLAST comparisons to characterized biosynthetic gene clusters. This resulted in the putative identification of 18 NRPS pathways for the biosynthesis of siderophores from the ALOHA gene catalog (Table S3). Combined, a total of 38 putative siderophore BGCs consisting of 101 unique genes were identified and used for downstream analysis (Table S3). The antiSMASH pipeline was utilized to assign putative structural characterizations to these siderophore BGCs based on comparisons to known biosynthetic pathways. We note that while this is a conservative approach to identifying siderophore BGCs that may exclude siderophores without well characterized biosynthetic pathways, it has resulted in a list of genes with high percent identity to known siderophore biosynthesis genes, giving us confidence in the functional annotation of genes used for downstream analysis.

To examine the distribution of siderophore BGCs with depth and across time at Station ALOHA, metagenomic sequence reads from plankton biomass (> 0.2 μm) collected on HOT cruises between May 2015 and April 2016 (NCBI BioProject PRJNA352737) were mapped to open reading frames (ORFs) of the ALOHA gene catalog. In total, this consisted of 132 samples from 5-500 m, spanning 11 months. Forward and reverse reads from each sample were quality trimmed with Trimmomatic (Bolger et al. 2014) and mapped to the ALOHA gene catalog using BWA (Li and Durbin 2009) with a 95% nucleotide identity threshold and minimum alignment length of 45 bp. Reads aligned as proper pairs were then counted with FeatureCounts (Liao et al. 2014) and multiple mappers were fractionally distributed. Finally, read counts were normalized to reads per kilobase per million (RPKM) for further analysis. Reads from the 2015-2016 dataset recruited to 24 of the 38 siderophore BGCs identified in the ALOHA gene catalog (Table S3).

## 3. Results

### 3.1 Uptake and turnover time of siderophore-bound iron by the microbial community

Uptake rates of inorganic Fe were the highest across all treatments (Fig. 1B, Table S1). The inorganic Fe treatments were denoted as FeL, since the added FeCl_3_ likely associated with unbound natural ligands present in excess of the in situ dFe in the experiment (Bundy et al. 2018). Uptake rates of FeL at both 15 m and 300 m were higher on average in the 0.2-2.0 µm size fraction compared to the >2.0 μm size fraction. In the >2.0 µm size fraction, uptake rates of FeL at 15 m were comparable in the light and the dark (∼36 pmol L^-1^ day^-1^). In contrast, FeL uptake was higher (on average) in the light (85 pmol L^-1^ day^-1^) compared to the dark (50 pmol L^-1^ day^-1^) in the 0.2-2.0 µm size fraction (Fig. 1B).

Siderophore-bound Fe was readily accessible by the microbial community at 15 m and 300 m at Station ALOHA (Fig. 1B, Table S1). The relative uptake rate varied for the different Fe-siderophore treatments. At both depths, the Fe-amphibactin amendment was taken up the fastest (54 pmol L^-1^ day^-1^), followed by ferrioxamine E (19 pmol L^-1^ day^-1^) and ferrioxamine B (9 pmol L^-1^ day^-1^). In contrast to FeL, the uptake rate of siderophore-bound Fe was generally higher in the dark (33 pmol L^-1^ day^-1^) compared to the light (14 pmol L^-1^ day^-1^). Similar to the FeL treatment, Fe uptake was higher on average in the 0.2-2.0 µm fraction compared to the >2.0 µm fraction across all siderophore treatments (Fig. 1B).

Surprisingly, the uptake rate of siderophore-bound Fe was higher at 300 m compared to 15 m leading to shorter Fe turnover times (Fig. 1C). In general, at both 15 m and 300 m the turnover times were shortest in FeL treatments, followed by Fe bound to amphibactins, ferrioxamine E and ferrioxamine B. The average turnover time of Fe at 15 m ranged from 15±8 days in the FeL treatment to 252±376 and 215±165 days in the ferrioxamine E and ferrioxamine B treatments, respectively. Mean turnover times at 300 m ranged from 9±4 days in the FeL treatment to 65±4 days in the ferrioxamine B treatment (Fig. 1C). Across all treatments, turnover times of Fe were consistently shorter at 300 m compared to 15 m (Fig. 1C), though this difference was only significant in the ferrioxamine B treatment (pairwise t-test, *p* < 0.05).

The Fe uptake rate per cell (amol Fe day^-1^ cell^-1^) was calculated from the total Fe uptake rate. The Fe uptake rates in the dark from the 0.2-2.0 and >2.0 µm size fractions were summed for a given Fe substrate (Table S1) and normalized to the abundance of heterotrophic bacteria measured via flow cytometry, assuming that the Fe uptake in the dark treatments was dominated by these organisms. This is likely a valid assumption for the experiments done at 300 m, but for experiments done at 15 m there could be additional Fe uptake in the dark by phytoplankton. The resulting Fe uptake per cell at 15 m ranged from 0.01-0.14 amol Fe cell^-1^ day^-1^, increasing to 0.18-1.46 amol Fe cell^-1^ day^-1^ at 300 m given the higher total rates of Fe uptake and much lower cell abundances (Table S1).

Cellular Fe:C ratios were calculated in two different ways (see section 2.2). The first, determined an Fe quota per cell (pmol Fe cell^-1^) at the end of each experiment assuming a bacterial growth rate of 1 day^-1^ (Jones et al. 1996), such that the Fe uptake rate per cell is equal to the cellular Fe quota. These cellular Fe quotas were then converted to Fe:C ratios using a fixed carbon quota of 12.4 fg C cell^-1^ (Strzepek et al. 2005; Boyd et al. 2015). The second method compared total Fe uptake rates (pmol Fe L^-1^ day^-1^) to average rates of bacterial carbon production measured at or near Station ALOHA (23 nmol C L^-1^ day^-1^ and 0.9 nmol C L^-1^ day^-1^ at 15 m and 300 m, respectively). This represents the ratio at which Fe and C are actively being acquired by bacterial cells and does not require assumptions of growth rates or cellular carbon quotas, but does presume a constant leucine:carbon ratio within biomass. At 15 m, the Fe:C per cell calculated using a fixed carbon quota ranged from 6.8±3.5 µmol:mol (ferrioxamine B treatment) to 68±43 µmol:mol (FeL treatment; Table S1). In contrast, the Fe:C calculated using average rates of bacterial production ranged from 380±192 µmol:mol to 3775±2386 µmol:mol (Table S1). We note that assuming a lower heterotrophic bacterial growth rate of 0.05 day^-1^ in surface waters (Kirchman 2016) with a fixed carbon quota results in higher Fe:C ratios determined by the first method, and would reduce the discrepancy between these two methods. We however chose to keep the estimates using a growth rate of 1 day^-1^ as our lower bound since these estimates are comparable to previous Fe:C values reported previously (Tortell et al. 1999). Irrespective of which approach was used, the Fe:C per cell was higher at 300 m than 15 m. The Fe:C per cell calculated from a fixed carbon ratio at 300 m ranged from 88±6 µmol:mol (ferrioxamine B treatment) to 711±305 µmol:mol (FeL) treatment, while Fe:C per cell based on bacterial production ranged from 17,044±1131 to 137,500±58,941 µmol:mol in the ferrioxamine B and FeL treatments, respectively (Table S1). These high Fe uptake rates were observed even after oxalate washing to remove adsorbed Fe and subtraction of a dead control, indicating that they do reflect internalization and uptake of Fe and not, for example, authigenic Fe precipitation.

### 3.2 Seasonal changes in nutrients and productivity at Station ALOHA during the study period

Station ALOHA is a well-studied oligotrophic time-series station in the North Pacific that is sampled by the HOT program on a nearly monthly basis (Fig. 1A). This region is characterized by low surface nutrient concentrations and a persistent deep chlorophyll maximum (DCM) centered around 100 m (Fig. 2A). During the study period, monthly mean nitrate+nitrite (NO_3_^-^ +NO_2_^-^) concentrations in the upper euphotic zone ranged from 1.4±0.4 nmol L^-1^ in August to 5.7±0.8 nmol L^-1^ in November (Fig. 2B, S1A). Below 100 m, NO_3_^-^+NO_2_^-^ concentrations increased rapidly and were more variable than the upper euphotic zone. Between 100-150 m, monthly mean NO_3_^-^+NO_2_^-^ concentrations during the study period reached minimum values in May (269.8±274.6 nmol L^-1^) and maximum values in August (1967.3±1858.3 nmol L^-1^, (Fig. 2B, S1B)). The unusually high mean NO_3_^-^+NO_2_ concentrations observed in August during the study period, along with the high degree of variability during this month, are likely the result of strongly negative sea level anomalies present at Station ALOHA during August 2015, increasing nutrient concentrations in the lower euphotic zone as a result of isopycnal uplift (Barone et al. 2019). Mean monthly depth integrated stocks of NO_3_^-^+NO_2_^-^ (mmol N m^-2^) in the upper and lower euphotic zone followed similar monthly patterns to that of average NO_3_^-^+NO_2_^-^ concentrations (Fig. S2).

**Figure 2.**
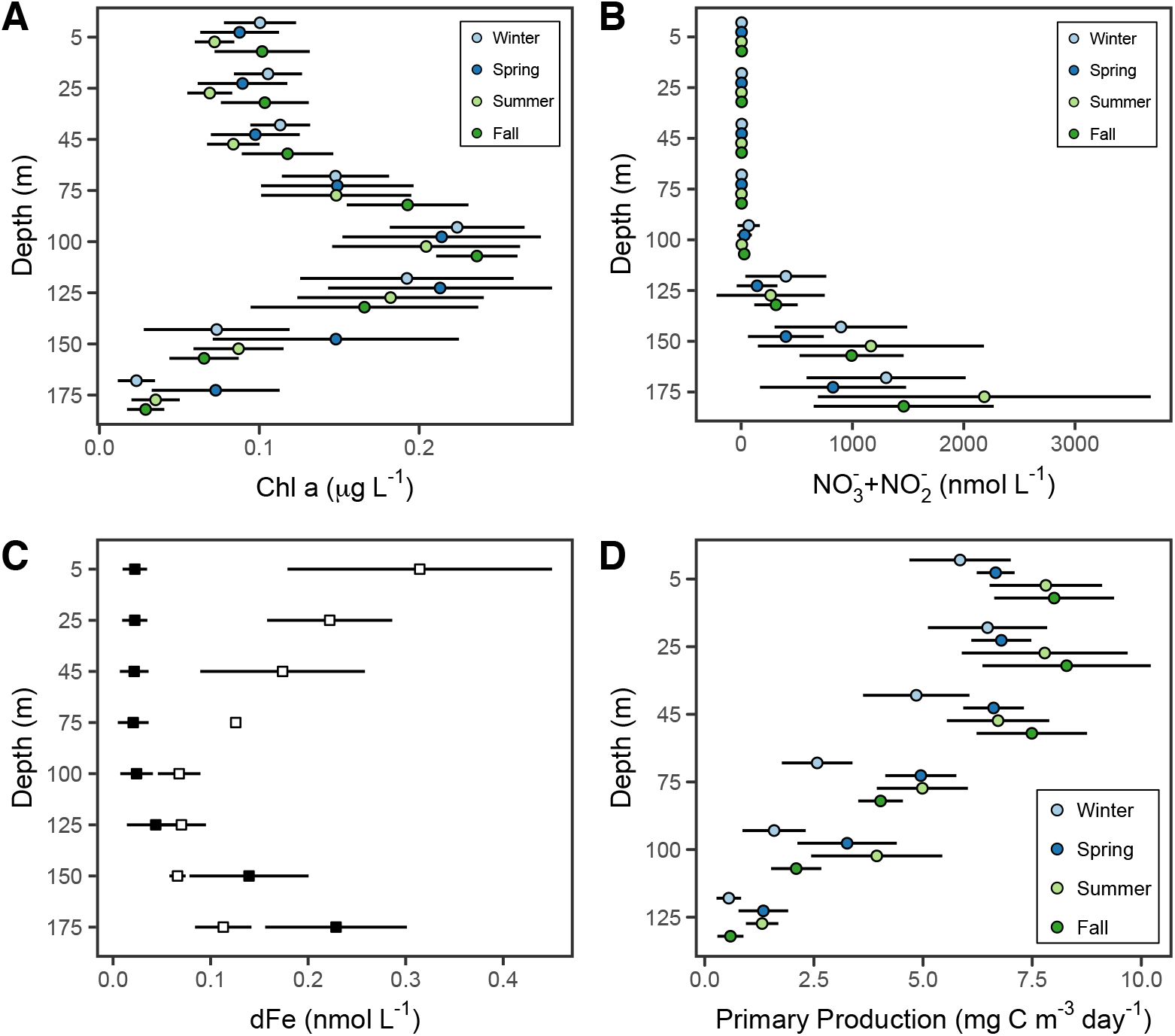
Depth profiles of **(A)** chlorophyll *a* (Chl *a*) concentrations (μg L^-1^), **(B)** nitrate+nitrite (NO_3_^-^+NO_2_^-^) concentrations (nmol L^-1^), **(C)** dissolved iron (dFe) concentrations (nmol L^-1^), and **(D)** rates of primary production (mg C m^-3^ day^-1^) during the 2013-2016 study period at Station ALOHA. Depth profiles in A, B, and C display the mean concentrations at each depth for a given season and error bars represent the standard deviation from the mean. Depth profiles in C display the mean dFe concentrations at each depth across the entire study period for available in situ measurements (open squares) and model output from the MIT Darwin model (closed squares) and error bars represent the standard deviation from the mean.

The dFe concentrations are persistently low in surface waters at Station ALOHA. Since limited dFe data was available during the study period, 3-day averaged dFe concentrations generated from the MIT-Darwin model were used as a potential means to explore seasonal variation in dFe concentrations, despite the known limitations of modeling dFe. Model output predicted minimum concentrations of dFe in the upper euphotic zone with low seasonal variability (Fig. 2C). However, dFe measurements collected during spring and summer months between 2012-2015 at Station ALOHA suggested dFe concentrations of the upper euphotic zone varied as much as 10-fold (0.08 to 0.87 nmol L^-1^), variability that is not captured in the model output (Fig. 2C). This high degree of variability in the upper euphotic zone is likely a result of episodic atmospheric dust deposition which can impact dFe concentrations in the upper euphotic zone (Fitzsimmons et al. 2015). Given the lack of agreement between model output and observations during the spring and summer in the upper euphotic zone, dFe availability in this region of the water column during fall and winter remains unconstrained without further measurements. Both model and observational data suggested consistently low (≤ 0.1 nmol L^-1^) dFe concentrations between 100-125 m, corresponding with the typical depth of the DCM. Below 125 m, dFe concentrations began to increase. However, observational data suggest there is a deeper ferricline than that indicated by model output.

During the study period, rates of primary production ranged from 0.2 – 11.8 mg C m^-3^ day^-1^ throughout the euphotic zone, with the lowest values observed in winter (Fig. 2D, S1A). Mean monthly rates in the upper euphotic zone peaked in October (7.8±2.4 mg C m^-3^ day^-1^) and were the lowest in February (4.8±1.9 mg C m^-3^ day^-1^; Fig. S1A). Between 100-150 m, mean monthly rates of primary production peaked in July (4.2±0.8 mg C m^-3^ day^-1^) with minimum values observed in December (1.1±0.3 mg C m^-3^ day^-1^; Fig, S1B). During the study period, depth integrated rates of primary production (0-150 m) increased from winter to summer before decreasing in fall (Fig. 3A). Maximum monthly mean values were observed in June (728.8±87.6 mg C m^-2^ day^-1^) and minimum values in December (453.4±23.0 mg C m^-2^ day^-1^). The particulate carbon flux at 150 m during the same time period showed similar monthly patterns to primary production (Fig. 3B), but with relatively higher fluxes observed during January and February. As a result, relatively small changes in the monthly mean *e*-ratio (calculated as the fraction of the total depth integrated rate of primary production from 0-150 m that is captured in the particulate carbon flux at 150 m) were detected, but followed similar patterns to those of particulate carbon flux (Fig. 3C). The highest monthly mean values were observed in February (0.062±0.031) and August (0.058±0.021) with the lowest in December (0.030±0.009). The average C:N ratio of the particulate flux at 150 m increased over the summer months from 6.8±0.3 in April to 8.4±1.2 in August, indicating that carbon-enriched particulate matter contributed to periods of high and relatively efficient export. However, the overall low *e*-ratios observed across the study period are consistent with intense remineralization of organic matter within the euphotic zone. Seasonal patterns in productivity observed during the 2013-2016 study period discussed here are consistent with those observed across the entire 30 year period of the HOT program (Karl et al. 2021).

**Figure 3.**
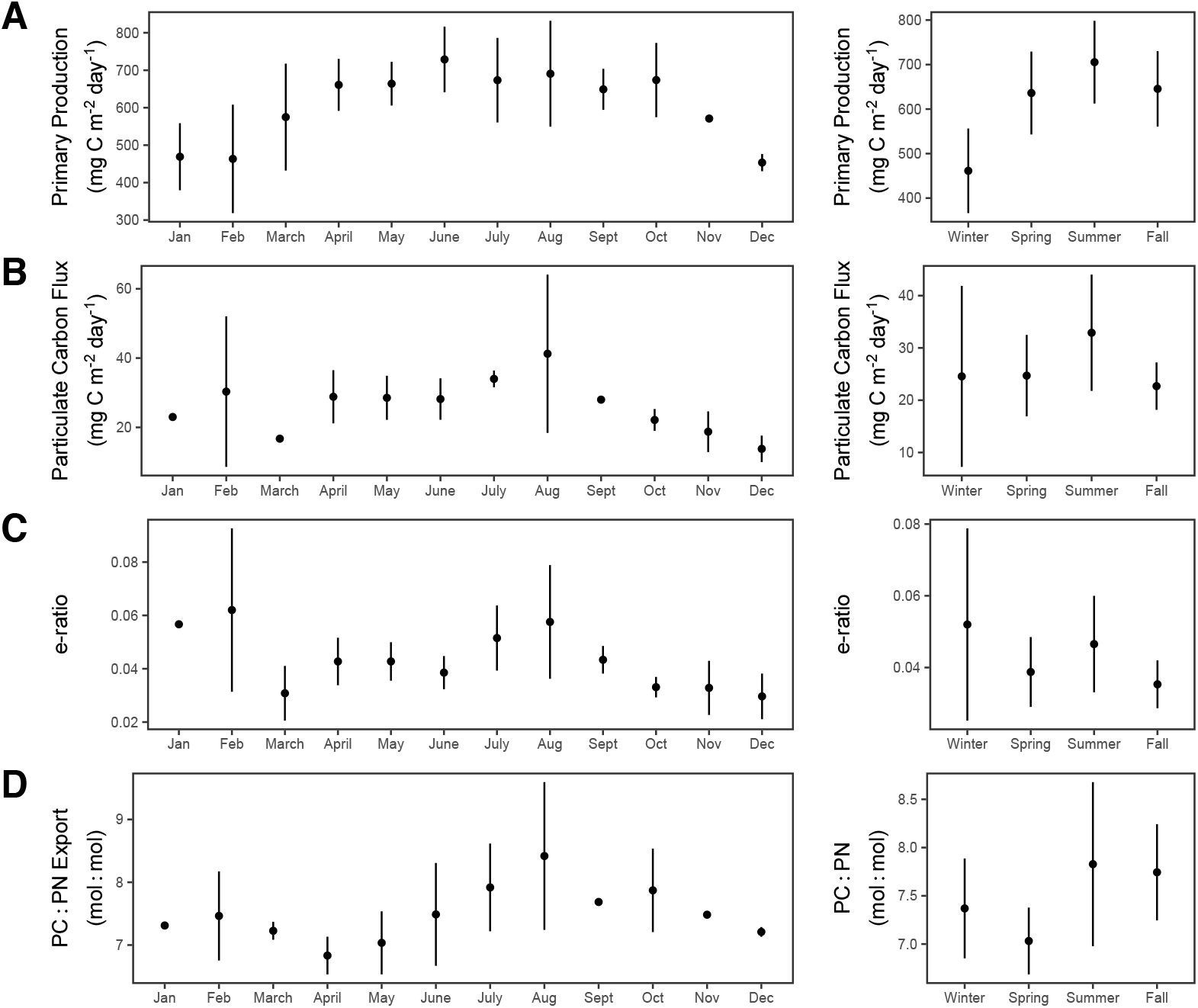
Monthly and seasonal mean values of **(A)** depth-integrated rates of primary production (mg C m^-2^ day^-1^) from 0-150 m, **(B)** particulate carbon flux (mg C m^-2^ day^-1^) at 150 m, **(C)** the *e*-ratio defined as the proportion of particulate carbon flux at 150 m compared to the 0-150 m depth-integrated rate of primary production, and **(D)** the C:N ratio of sinking particulate matter collected at 150 m at Station ALOHA during the 2013-2016 study period. Error bars represent the standard deviation from the mean.

### 3.3 Seasonal distribution of siderophores and siderophore biosynthesis genes at Station ALOHA

Dissolved siderophore concentrations and the abundance of siderophore BGCs changed seasonally at Station ALOHA as well as with depth (Fig. 4 and 5). Total dissolved siderophore concentrations varied from undetectable to 95.3 pmol L^-1^ during the entire study period. Note that in the fall, siderophore concentrations were only measured at 15 m, limiting assessment of seasonality in siderophore concentrations in the mesopelagic waters during this season. Average siderophore concentrations throughout the whole water column were highest in the spring (21.8±41.5 pmol L^-1^), fall (8.4±8.1 pmol L^-1^), and summer (3.0±4.2 pmol L^-1^) and lower in winter (0.4±0.6 pmol L^-1^, Fig. 5). However, high variability was observed within each season. In the upper euphotic zone where the most samples were collected, siderophore concentrations were highest on average in the spring (47.8±67.2 pmol L^-1^), fall (8.4±8.1 pmol L^-1^), and summer (2.4±0.1 pmol L^-1^), followed by winter (0.4±0.3 pmol L^-1^; Fig. 4, Table S2).

**Figure 4.**
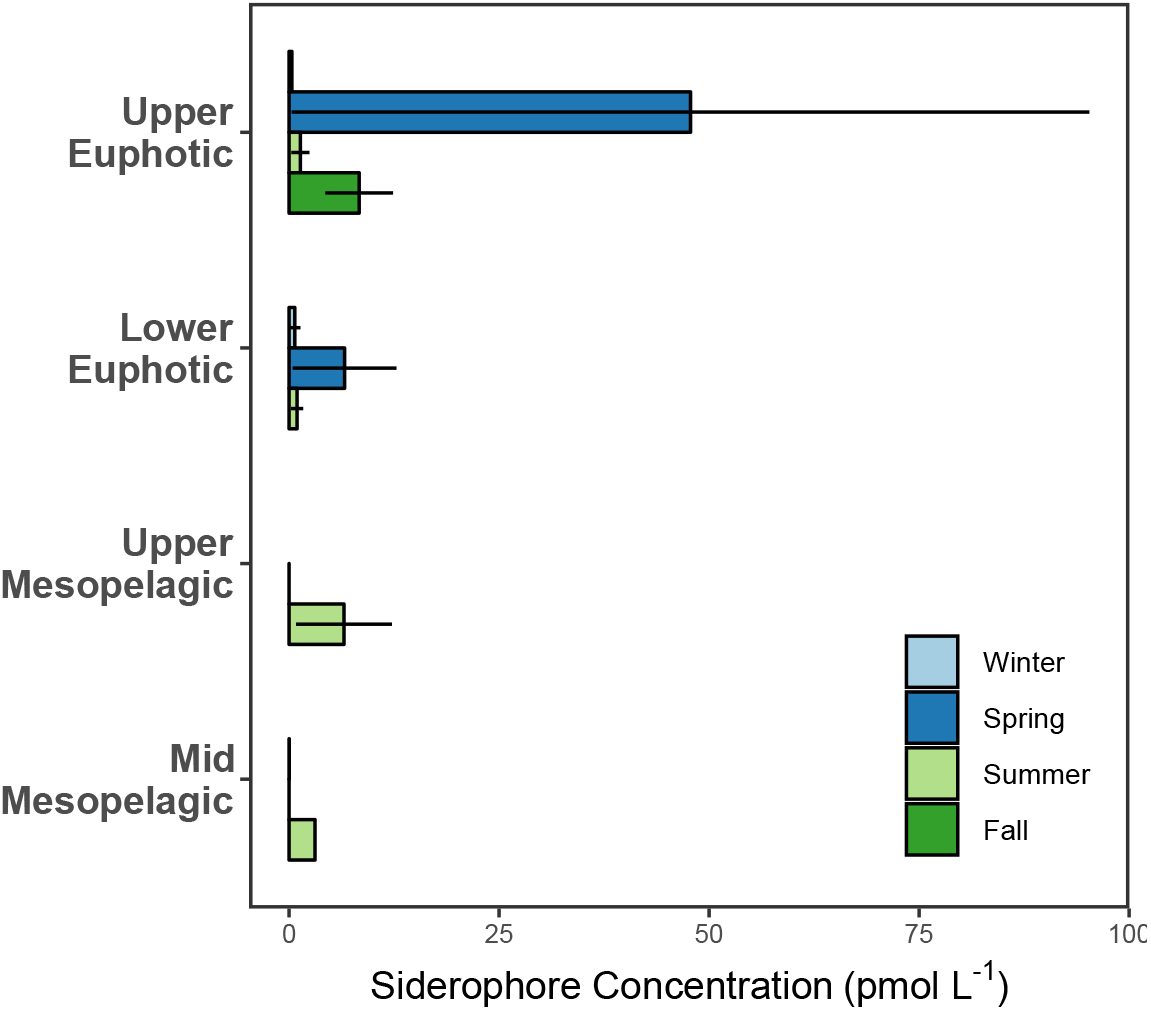
Seasonality in dissolved siderophore concentrations (pmol L^-1^) at Station ALOHA between 2013-2016. Bars display the mean values of measurements collected within the same depth range for a given season and error bars, when present, represent the standard error of the mean. When no data are present for a given depth range and season, no samples were collected during that time. Depth ranges are as follows: upper euphotic 0-75 m, lower euphotic > 75 m ≤ 150 m, upper mesopelagic > 150 m ≤ 300 m, mid-mesopelagic > 300 m ≤ 500 m.

**Figure 5.**
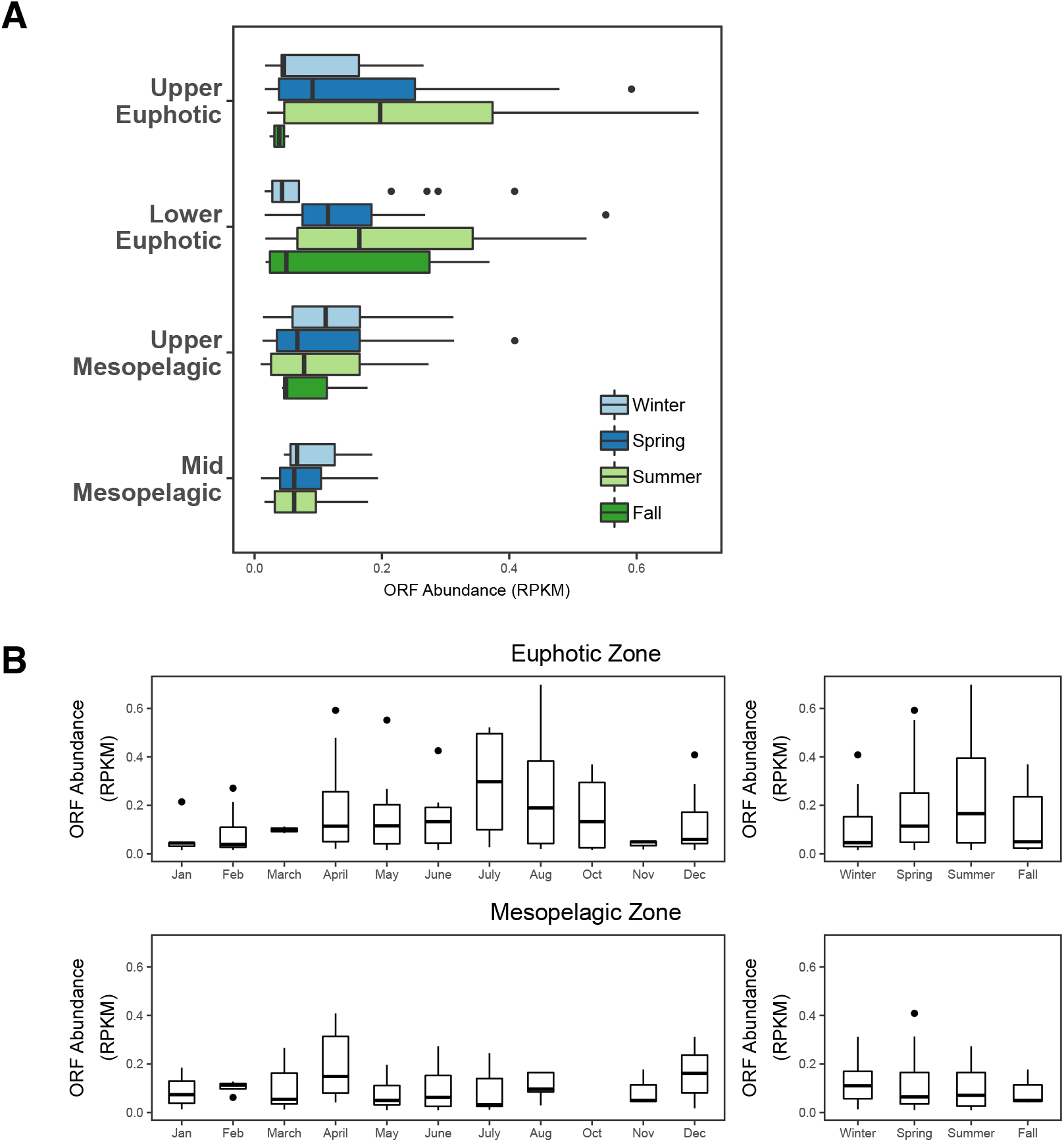
Seasonal abundance of ORFs (RPKM) from siderophore BGCs detected at Station ALOHA between May 2015-April 2016. **(A)** Box plots display the interquartile range of ORF abundances detected within a given depth range and season. Median values are plotted as solid vertical lines, and outliers are plotted as distinct points. Depth ranges are as follows: upper euphotic 0-75 m, lower euphotic > 75 m ≤ 150 m, upper mesopelagic > 150 m ≤ 300 m, mid-mesopelagic > 300 m ≤ 500 m. **(B)** Box plots display the interquartile range of ORF abundances detected within a given month or season in the euphotic (0-150 m) and mesopelagic (150-500 m) zones. Median values are displayed as solid horizontal lines and outliers are plotted as distinct points. No data available for September.

The capacity for siderophore production as indicated by the presence of ORFs from siderophore BGCs was detected at Station ALOHA across all seasons and at all depths sampled during the study period (Fig. 5A, Table S3). Similar to dissolved siderophore concentrations, ORFs from siderophore BGCs also showed low and relatively uniform median abundances (< 0.2 RPKM) in the winter in both the euphotic and mesopelagic zones, increasing in the euphotic zone in the spring (Fig. 5B). Monthly median ORF abundances in the euphotic zone peaked during the summer (July, 0.30 RPKM), decreasing throughout the fall in the upper euphotic zone, but remaining elevated in the lower euphotic zone. Lowest monthly median ORF abundances in the euphotic zone were observed in January and February (0.04 RPKM). Mesopelagic ORF abundances were less seasonally variable (Fig. 5B), with median ORF abundances ranging 0.05-0.11 RPKM. Monthly median ORF abundances were maximal in April (0.15 RPKM) and ORFs from siderophore BGCs were not detected in October in the mesopelagic. Several different siderophores were detected both as dissolved siderophores in the water column and as a BGC in the Station ALOHA gene catalog (Table 1). Of those identified in the water column, primarily hydroxamate siderophores were detected, with ferrioxamines and amphibactins being the most common (Table 1). The detected siderophore BGCs largely belonged to catecholate and carboxylate-type siderophores (Table 1). Two ferrioxamine BGCs were identified in the ALOHA gene catalog, however, these BGCs were not detected in the 2015-2016 metagenomic dataset. Several siderophores were found both in the water column and in the metagenomes, including petrobactin, pseudoalterobactin, piscibactin and vibrioferrin (Table 1). The biosynthesis genes were primarily homologues of known genes from *Vibrio*, *Photobacterium*, *Alteromonas*, and *Pseudoalteromonas* (Table S3).

**Table 1.**
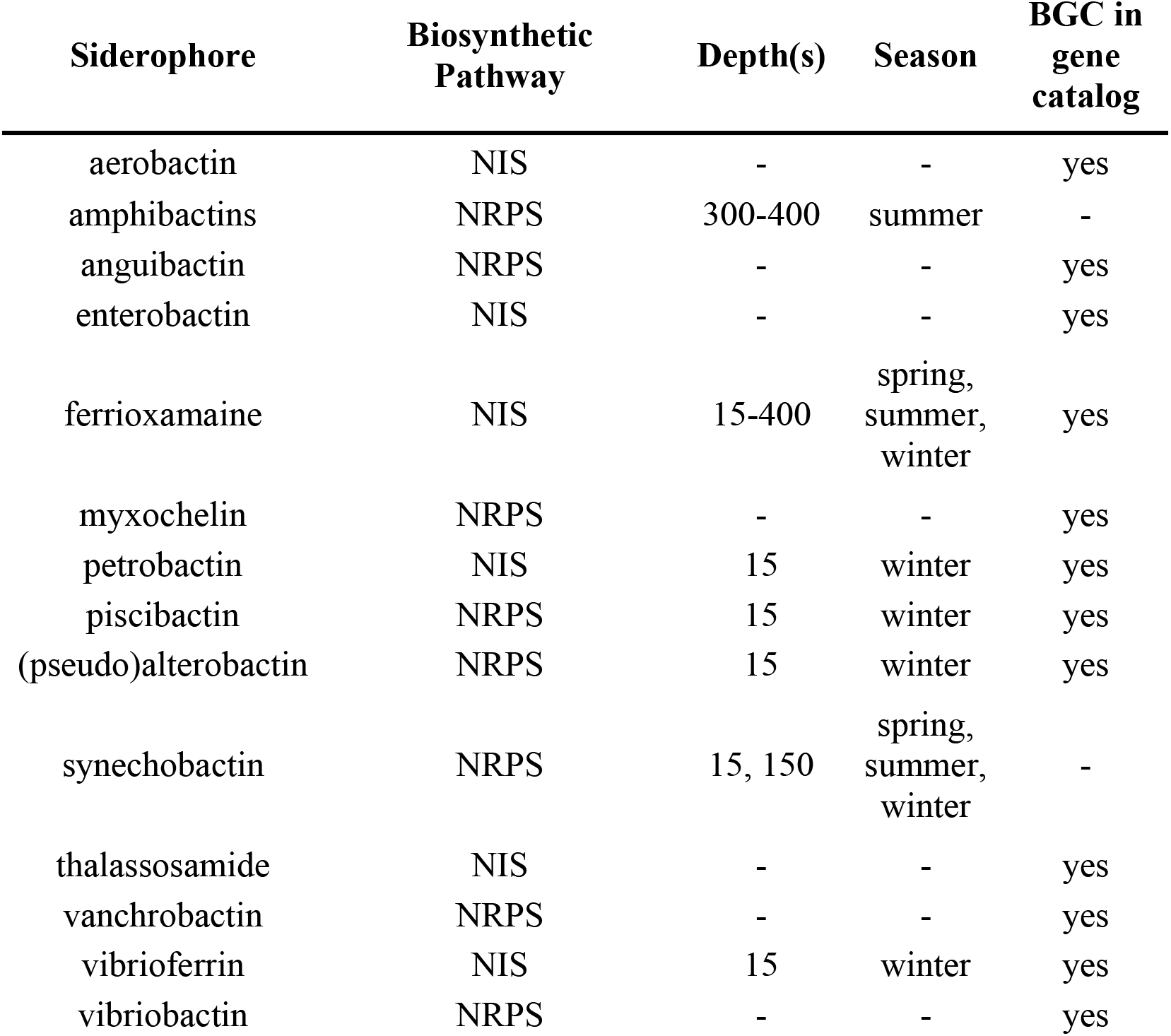
Siderophores measured in the water column and siderophore biosynthetic gene clusters (BGCs) identified at Station ALOHA. Siderophores identified in the water column are grouped according to type and are shown in the table based on that grouping. For example, multiple different forms of amphibactins were detected (Table S2) but are all grouped here under “amphibactin.” Specific siderophore IDs can be found in Supplementary Table 2. The ‘-’ denotes that the specific siderophore was not found either in the water column or in the Station ALOHA gene catalog.

## 4. Discussion

### 4.1 Turnover times of inorganic iron and iron-siderophore complexes reflect bioavailability to the microbial community

Siderophore mediated acquisition of dFe involves secretion, uptake, and recycling of these compounds, and the turnover time of each Fe source reflects its bioavailability. Siderophore-bound Fe was taken up less rapidly than the inorganic Fe additions, which were likely complexed by unbound ambient ligands. Siderophore Fe uptake had average turnover times ranging from 15-250 days (Fig. 1C) and an average turnover time of 148 days from all depths and siderophore treatments, while FeL turned over more rapidly averaging 14 days (Fig. 1C). The different turnover times reflects differences in the bioavailability of inorganic versus organically-bound Fe (Lis et al. 2015). The average turnover times of FeL observed in this study were very similar to turnover times of 14-18 days in the mixed layer during June 2019, measured at or near background dFe concentrations at Station ALOHA (Hawco et al. 2022). The shorter turnover of FeL likely reflects weaker binding of inorganic Fe associated with ligands naturally present in seawater, which are more accessible to phytoplankton using ferric reductases as an uptake mechanism (Maldonado and Price 2001; Salmon et al. 2006; Lis et al. 2015; Coale et al. 2019). Moreover, we found rates of FeL uptake were higher in the light than in the dark, suggesting significant uptake by photoautotrophs. Uptake rates of siderophore-bound Fe, in contrast, were always higher in the dark than the light, suggesting that siderophore Fe consumption was likely controlled by heterotrophic plankton and that photochemical impacts on the uptake of siderophore-bound Fe were minimal (Barbeau 2006). The turnover time of Fe bound to siderophores was also much longer, on the order of weeks to months, suggesting that Fe bound to siderophores is longer-lived than the inorganic and/or weakly bound Fe pool, and may help to retain dFe in both euphotic and upper mesopelagic waters.

Turnover times of siderophore-bound Fe varied depending on the siderophore. Fe associated with amphibactins was taken up more quickly than ferrioxamine E or B (Fig. 1B). Amphibactins have a lower binding affinity to dFe (log of the conditional stability constants = 12.0-12.5; Bundy et al. 2018) compared to ferrioxamines (log of the conditional stability constants = 14.0-14.4; (Bundy et al. 2018)), and therefore Fe could be more easily reducible from amphibactins. Alternatively, more members of the microbial community may possess direct Fe-amphibactin uptake pathways, but these uptake pathways are not well constrained. The relative turnover times and bioavailability of the different siderophores therefore, appeared to be governed by their binding strengths in this case. In our experiments, the highest uptake rates of siderophore-bound Fe were also found in the 0.2-2.0 μm size fraction and in the dark, suggesting Fe bound to siderophores may be more accessible to heterotrophic bacteria rather than larger photoautotrophs.

### 4.2 Rates of iron acquisition by the microbial community are elevated in the mesopelagic waters of Station ALOHA relative to the surface

An unexpected result of this work was the faster uptake rates and shorter turnover times of Fe observed at 300 m compared to 15 m, especially in the case of siderophore-bound Fe, resulting in higher uptake rates per cell and higher Fe:C ratios at depth. There are several experimental details that should be considered when interpreting these observations. While these factors were controlled for whenever feasible, the possibility remains that they have influenced the results presented here. For example, macronutrients are elevated at 300 m relative to 15 m, so it is possible Fe uptake rates were enhanced due to relief from macronutrient limitation. However, our surface water treatments included modest NO_3_^-^ amendments that should have alleviated nitrogen limitation. The experiments at 300 m were also incubated at the same temperature as the 15 m experiments, likely enhancing the uptake rate of Fe relative to *in situ* rates due to increased metabolic activity at higher temperatures. However, it is not clear why this would stimulate uptake rates above those measured in surface waters. An additional possibility is that the ^55^Fe adsorbed to the outside of cells or detritus more readily at 300 m, causing apparent high Fe uptake compared to 15 m. If this was the case, we would have expected the dead controls from 300 m to have significantly higher activities than those from 15 m, and this was not observed. Furthermore, oxalate rinsing was applied to all samples, which should remove most (if not all) adsorbed Fe. Finally, in terms of the observed Fe:C ratios, an additional source of uncertainty may come from the conversion of Fe uptake to Fe:C using a fixed carbon quota. Fe:C ratios based on a fixed carbon quota overlap with published values for surface waters, but resulted in values for 300 m that were significantly higher (Table S1). If the actual carbon quota per cell is significantly higher in heterotrophic bacteria at 300 m, then the calculated Fe:C ratios presented here could be artificially elevated. Carbon biomass estimates are not available for any cultured marine bacteria from mesopelagic waters, so we are not able to determine if this fixed carbon quota is too low.

Accounting for these possible experimental artifacts, it is likely that the fast uptake rates and short turnover times of Fe observed at 300 m are related to ecologically relevant differences in Fe acquisition by the microbial community at this depth. The Fe requirements of heterotrophic bacteria are not well known, but evidence suggests they are elevated relative to large phytoplankton (Fe:C ∼ 2.0 µmol: mol) or cyanobacteria (Fe:C ∼ 19 µmol: mol) (Strzepek et al. 2005). Published estimates for heterotrophic bacteria Fe:C ratios range from 7.5 (Tortell et al. 1996) to 83 µmol: mol (Mazzotta et al. 2020), and the observations at 300 m in this study expand that range to upwards of 700 µmol: mol. The range of Fe requirements observed across these studies likely varies according to experimental growth conditions and the lifestyle strategies of specific groups of bacteria under consideration. In the current study, the 1 nmol L^-1^ dFe additions and slightly higher in situ dFe concentrations at 300 m may have supported luxury Fe uptake and storage by the *in situ* bacteria community. This could lead to higher Fe uptake at these depths, and much higher Fe:C quotas than for surface bacteria communities. Previous work has identified multiple copies of the Fe-storage protein bacterioferritin in a cultured marine heterotroph (Mazzotta et al. 2020) and modeling work suggests that bacteria competitively consume Fe and that Fe storage plays a large role (Ratnarajah et al. 2021). Alternatively, it is possible that the consumption of refractory dissolved organic matter at these depths has a higher Fe requirement for marine bacteria, though this has not been explored to our knowledge. The energy starved microbial community in the upper mesopelagic could have elevated metabolic requirements for Fe (Tortell et al. 1996) relative to the surface ocean community, potentially resulting from differences in the lability or oxidation state of organic matter. Additionally, nitrifying bacteria and archaea, whose cellular abundances significantly increase in mesopelagic waters, appear to have high Fe requirements (Shafiee et al. 2019; Saito et al. 2020). In addition to the high uptake rates and Fe:C quotas observed here, other lines of evidence are starting to point to an increased demand for Fe in the mesopelagic. For example, high concentrations of siderophores have been observed in the upper mesopelagic ocean, suggesting active Fe uptake at these depths (Bundy et al. 2018; Park et al. 2023; Li et al. 2024). Ultimately, understanding the Fe requirements of mesopelagic bacteria could have important implications for carbon and Fe cycling in the twilight zone.

### 4.3 Siderophore production at Station ALOHA reflects seasonal variation in primary production and organic matter availability

Measurements of dissolved siderophores highlight seasonal differences at Station ALOHA, with siderophore concentrations greatest in the spring, summer and fall but almost entirely absent or low in winter (Fig. 4). There was high variability in total dissolved siderophore concentrations and the identity of siderophores observed over the study period. Some of this variability may have been caused by factors related to method developments over time. For example, siderophore concentrations were relatively high during our only fall sampling period (2013), but we were not able to identify specific siderophores with high confidence in these samples (Table S2). These samples were collected during the development of our LC-ICP-MS and LC-ESI-MS methods, and since that sample collection, continued improvements in detection limits and ionization parameters have enhanced our ability to detect siderophores with high confidence. Collecting profile samples during this time was not possible, because of the high volumes of water that were needed (800 L). However, improvements to methodological sensitivity prior to sample collection in 2014 allowed for the high confidence detection of several siderophores from the winter cruise in this year despite the relatively low concentrations that were observed (Table S2). The dissolved siderophore distributions were notably higher in the spring, summer, and fall, even when only considering surface waters (Fig. 4). The elevated concentrations of dissolved siderophores in spring and summer were matched by a higher potential for the microbial community to biosynthesize siderophores during these seasons, particularly in the euphotic zone (Fig. 5). Of the 14 different types of siderophores identified using mass spectrometry or from gene pathways in the metagenomes, 5 were found in both (Table 1). The remaining compounds were either only found in the water column and not in the metagenomes (2 of the 14), or conversely, the biosynthesis genes were found (7 of the 14) but the compound was not identified in the water column samples. A wide structural diversity of siderophores has been detected using the LC-MS techniques employed here (Boiteau et al. 2019), making it unlikely that specific siderophores escaped detection in the water column due to methodological bias. Therefore, the limited overlap in the dominant siderophores measured and the BGCs identified suggests that either some compounds may be underestimated from metagenomic approaches, or that some siderophores cycle too rapidly to be observed in measurable quantities. Further uptake studies examining the bioavailability of the siderophore groups dominant within the metagenomes, such as catecholate-type siderophores, will help to further explain these differences.

Seasonality in nutrient supply, productivity, and export have all been documented in the North Pacific subtropical gyre and Station ALOHA (Fitzsimmons et al. 2015; Hawco et al. 2021; Karl et al. 2021) and all impact microbial community structure and thus siderophore production. In the spring, primary production begins to increase at Station ALOHA (Fig. 2 and 3). This is thought to be a result of increasing solar irradiance (Karl et al. 2021) and these changes are especially evident in the lower euphotic zone ((Letelier et al. 2004, 2017); Fig. S1 and S2). This springtime increase in productivity generally coincides with the peak in Fe delivery to this region (Fitzsimmons et al. 2015). During this time, Fe is delivered primarily via atmospheric dust deposition which introduces lithogenic Fe particles to the surface ocean. In this study, the highest concentrations of dissolved siderophores were observed in surface waters during the spring and coincided with increasing biosynthetic capacity in the euphotic zone (Fig. 4 and 5). It is likely that siderophore secretion by heterotrophic bacteria at Station ALOHA during this time may help solubilize particulate Fe (Bundy et al. 2018) and to compete with other microorganisms for Fe (Boiteau et al. 2016) as the Fe demand of the microbial community begins to increase during these productive months. For example, ferrioxamines have been shown to be produced during the degradation of sinking (Velasquez et al. 2016) or suspended particles (Bundy et al. 2018), and high siderophore concentrations were observed in the North Pacific Transition Zone in a region of high dust inputs (Park et al. 2023).

As primary production increases throughout the summer months at Station ALOHA, essential nutrients such as NO_3_^-^ and dFe are consumed. During late summer, when the physical delivery of NO_3_^-^ to the surface ocean is low, the nitrogen fueling these higher rates of production is thought to be supplied by N_2_ fixation (Karl et al. 2021). Rates of N_2_ fixation peak in the late summer and early fall at Station ALOHA when warm, stratified waters select for the growth of N_2_-fixing cyanobacteria (Böttjer et al. 2017). This late summer production is also associated with a large pulse of particulate carbon export that reaches the sea floor and is thought to be the result of diatom growth supported by N_2_-fixing endosymbionts (Karl et al. 2012). As a result of increasing primary production and temperature, rates of heterotrophic bacterial production are also elevated by late summer at Station ALOHA (Viviani and Church 2017). However, during this period, dust delivery to the surface ocean decreases (Fitzsimmons et al. 2015). Thus, an increased Fe demand of the microbial community resulting from high rates of primary production, N_2_ fixation, and bacterial production coincides with low rates of Fe delivery and likely intensifies the competition for Fe throughout the euphotic zone as summer progresses. This has the potential to set up conditions favoring the production of siderophores by bacteria as a means to competitively acquire Fe or to retain or recycle Fe in the euphotic zone over longer timescales (Boyd et al. 2015; Hayes et al. 2015; Hawco et al. 2022).

In addition to dissolved siderophore concentrations, the genetic potential for siderophore biosynthesis also peaked during the summer months. Furthermore, a majority of the genera found to have siderophore BGCs in this study, such as *Vibrio* and *Alteromonas*, contain known copiotrophic bacteria, suggesting that siderophore production may be common to heterotrophs responding to episodic inputs of organic matter during periods of elevated production (Fontanez et al. 2015; Pelve et al. 2017; Church et al. 2021; Poff et al. 2021; Leu et al. 2022). These genera have been found to be dominant in sediment trap material from Station ALOHA and in association with sinking eukaryotes (Fontanez et al. 2015). Thus in many cases, siderophore production might largely be due to the Fe demand of heterotrophic bacteria as they consume carbon from sinking particles (Hopkinson and Barbeau 2012). In this study, seasonal patterns in the genetic potential for siderophore production within the euphotic zone closely followed those of the abundance of both suspended and sinking particles, further suggesting that siderophore biosynthesis may be directly tied to the dynamics of particle production. While the genetic potential for siderophore production did not vary seasonally in the mesopelagic zone, elevated dissolved siderophore concentrations were observed in the summer in this region of the water column, potentially in response to the elevated particle flux escaping the euphotic zone during this time. Finally, in winter, siderophore concentrations and the abundance of siderophore BGCs were reduced significantly throughout the water column, likely reflecting the decrease in metabolic activity or Fe demand of heterotrophic bacteria as productivity and the availability of organic substrates diminish, or increased mixing that dilutes siderophores.

Together, the higher concentrations of siderophores from spring to fall and the dominance of copiotrophic bacteria identified as potential siderophore producers in the metagenomes, predict that siderophore production is heightened during periods of enhanced productivity at Station ALOHA and provides compelling evidence that ligand production by the microbial community plays an important role in the seasonal cycling and retention of Fe at Station ALOHA. Uptake rates of Fe in the euphotic zone agree with previous measurements at Station ALOHA and imply that Fe is cycled rapidly by the microbial community in this region. While less bioavailable than inorganic Fe additions, siderophore-bound Fe proved to be a bioavailable source of Fe to the microbial community in both the euphotic and mesopelagic zones. Importantly, we found the resulting bioavailability depended on the specific siderophore structure, warranting additional studies focused on elucidating specific compounds in the Fe-binding ligand pool and their uptake kinetics in natural communities. Due to longer turnover times of Fe bound to siderophores, the production of strong siderophores has the potential to transform dFe into a form that persists in the water column on longer time scales. The finding that Fe uptake rates and turnover times were faster in the mesopelagic waters than in the surface ocean was surprising and merits further investigation. Such results suggest Fe availability is impacting microbial dynamics and Fe cycling at depth.

## 5. Acknowledgements

Special thanks to the Captain and crew of the R/V *Kilo Moana* and R/V *Ka’imikai-O-Kanaloa* for help with sample collection, to the Hawaii Ocean Time-series program for the numerous biogeochemical measurements and to Kendra A. Turk-Kubo for additional flow cytometry analyses. We would also like to thank Travis Mellett for helpful comments on the manuscript. RMB was funded by a Life Sciences-Simons Early Career Investigator in Marine Microbial Ecology and Evolution Awards (Award #618401) and a Woods Hole Oceanographic Institution postdoctoral fellowship. LEM was funded by a Simons Postdoctoral Fellowship in Marine Microbial Ecology (Award #729162). MJC was funded by the Simons Collaboration on Ocean Processes (Award #721221). NJH was supported by Life Sciences-Simons Early Career Investigator in Marine Microbial Ecology and Evolution Awards (Award #924096). The authors have no conflict of interest to declare.

## Data Availability Statement

Iron uptake data are available as supplementary files and on Zenodo (DOI: 10.5281/zenodo.7062571). Sequence data are available from the NCBI short read archive (SRA) under Bioproject no. <u>PRJNA352737</u>.

## Supplementary Information

**Table S1.** All ^55^Fe uptake data from each experiment. The treatments are ^55^Fe-amphibactins (Amp), ^55^Fe-ferrioxamine B (DFB), ^55^Fe-ferrioxamine E (DFE) and ^55^FeCl_3_ (FeL).The Fe:C^+^ of heterotrophic bacteria was calculated by assuming all dark treatments were dominated by heterotrophic bacteria Fe uptake, so the total uptake (sum of 2.0 + 0.2 µm uptake) in the 250 mL bottle over the course of 12 hours was converted to total moles of Fe and then divided by the total bacteria abundance at the end of the experiment determined by flow cytometry. A bacteria carbon per cell of 12.4 fg C cell^-1^ (Boyd et al. 2015) was used to calculate bacteria carbon from each treatment. The additional column of Fe:C* were calculated based on the comparison of iron uptake rates (pmol Fe L^-1^ day^-1^) to rates of bacterial carbon productivity (nmol C L^-1^ day^-1^) at Station ALOHA at 15 m and 300 m depth.

**Table S2.** Dissolved siderophore concentrations (pM) and identifications from all six cruises in different seasons. Only siderophores that were found complexed to iron are presented. The ‘-‘ means that a high confidence (confirmed with MS^2^) siderophore identification was not possible in that sample.

**Table S3.** The complete list of siderophore biosynthetic genes identified in the ALOHA gene catalog with functional and taxonomic identifications given where possible. Genes specifically identified in the 2015-2016 metagenomic dataset collected at Station ALOHA are noted as such.

**Figure S1:**
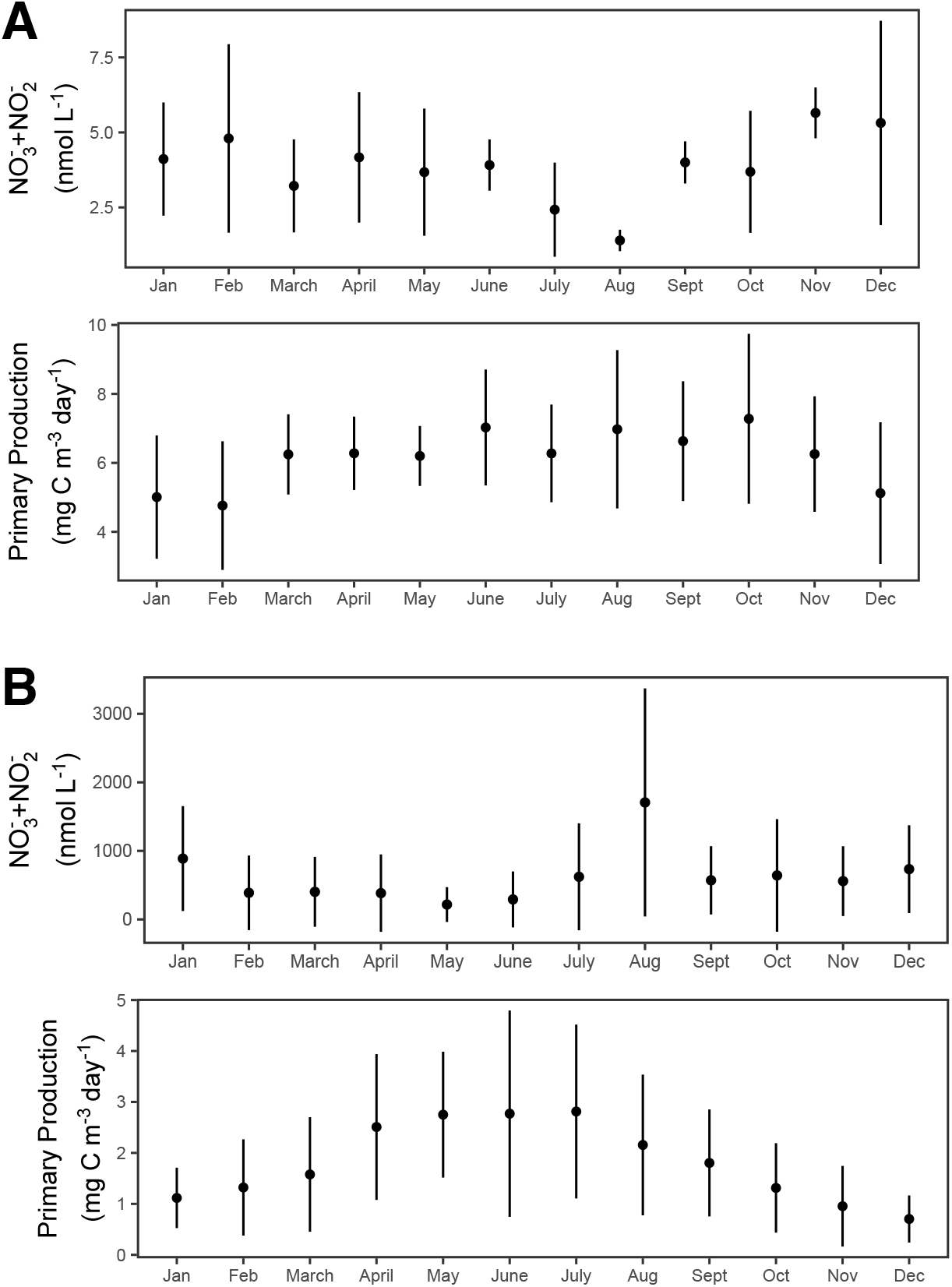
Monthly mean nitrate+nitrite (NO_3_^-^+NO_2_^-^) concentrations and rates of primary production in the **(A)** upper euphotic zone (≤ 75 m) and **(B)** lower euphotic zone (> 75 ≤ 150 m) at Station ALOHA during the 2013-2016 study period.

**Figure S2:**
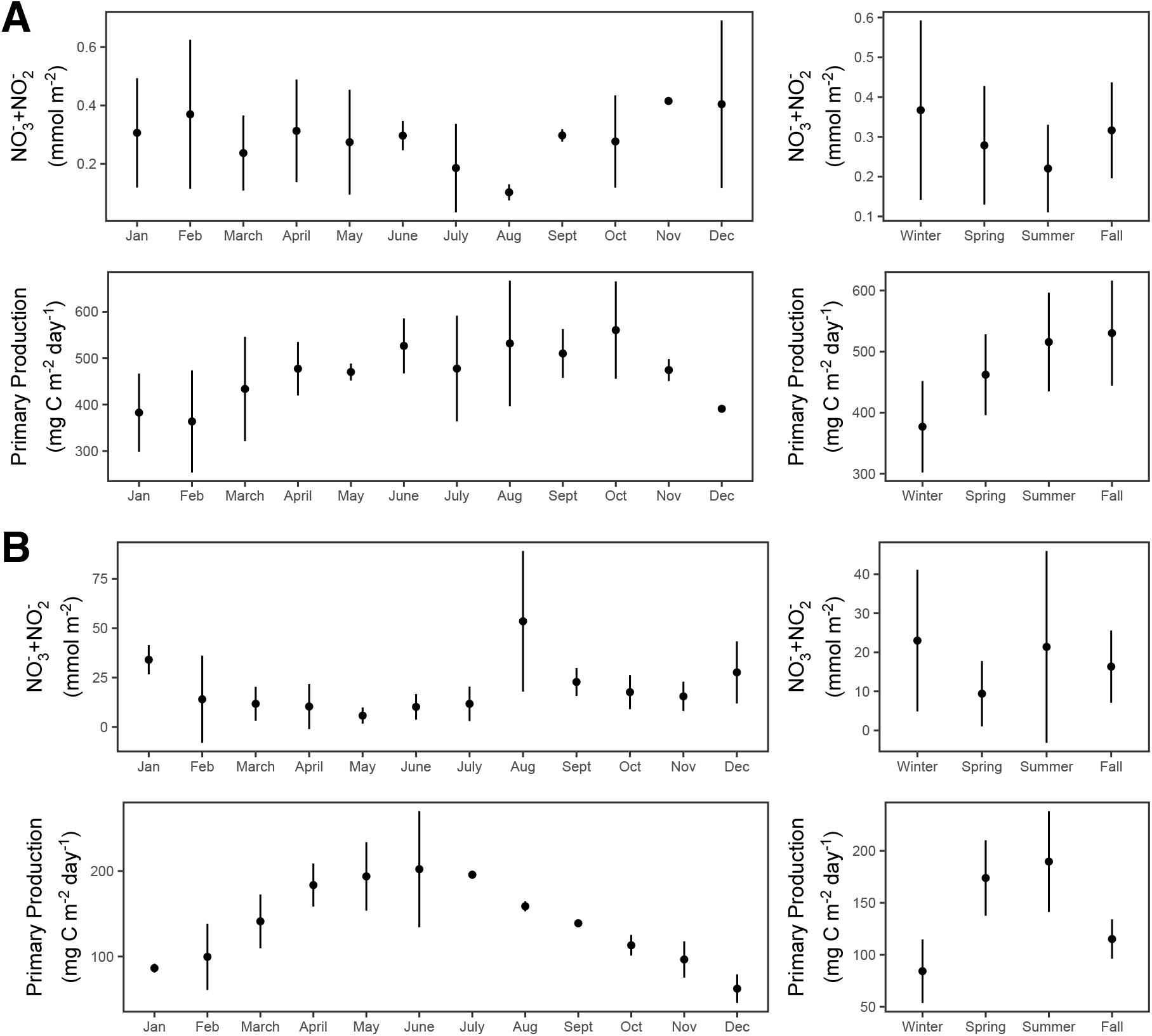
Monthly and seasonal mean depth-integrated concentrations of nitrate+nitrite (NO_3_^-^ +NO_2_^-^) and rates of primary production in the **(A)** upper euphotic zone (≤ 75 m) and **(B)** lower euphotic zone (> 75 ≤ 150 m) at Station ALOHA during the 2013-2016 study period.

